# A metal ion orients mRNA to ensure accurate 2’-*O* ribosyl methylation of the first nucleotide of the SARS-CoV-2 genome

**DOI:** 10.1101/2021.03.12.435174

**Authors:** Thiruselvam Viswanathan, Anurag Misra, Siu-Hong Chan, Shan Qi, Nan Dai, Shailee Arya, Luis Martinez-Sobrido, Yogesh K. Gupta

## Abstract

The SARS-CoV-2 nsp16/nsp10 enzyme complex modifies the 2’-OH of the first transcribed nucleotide of the viral mRNA by covalently attaching a methyl group to it. The 2’-*O* methylation of the first nucleotide converts the status of mRNA cap from Cap-0 to Cap-1, and thus, helps the virus evade immune surveillance in the host cell. Here, we report two structures of nsp16/nsp10 representing pre- and post-release states of the RNA product (Cap-1). We observe overall widening of the enzyme upon product formation, and an inward twisting motion in the substrate binding region upon product release. These conformational changes reset the enzyme for the next round of catalysis. The structures also identify a unique binding mode and the importance of a divalent metal ion for 2’-*O* methylation. We also describe underlying structural basis for the perturbed enzymatic activity of a clinical variant of SARS-CoV-2, and a previous SARS-CoV outbreak strain.

## Main

RNA viruses employ a diverse set of protein assemblies to enzymatically modify the 5’-end of their genomes. This process, termed RNA capping is essential for efficient a) production of viral proteins, b) protection of viral (v)RNA from host degradation, and c) subversion of the host innate immune responses – all of which enable viruses to thrive inside the host body^1^. In coronaviruses (CoVs), the nonstructural protein 16 (nsp16), and its non-catalytic stimulator nsp10, assemble on the 5’-end of nascent mRNA to perform the last step of RNA cap modification – *S*-adenosyl-L-methionine (SAM)-dependent methylation of the 2’-OH on the first transcribed nucleotide base (N_1_), usually an adenine. This converts the status of RNA from Cap-0 (^me7^Gppp**<u>A</u>**) to Cap-1 (^me7^GpppA**m**)^2,3^. Suppression of innate host antiviral response by Cap-1 through IFIT (interferon signaling-induced proteins with tricopeptide repeat) proteins^4^ is thought to be a consequence of diminished binding of IFIT to Cap-1^5^. Genetic ablation of nsp16 enzymatic activity also leads to induction of type I interferon (IFN) via the RNA sensor melanoma differentiation-associated protein 5 (MDA5)^6^. Emerging evidence suggests that COVID-19 patients have elevated innate immune responses, causing hypercytokinemia^7^. Thus, structural elucidation of different stages of Cap-1 formation and modification will further our understanding of the 5’-RNA biology of CoVs, and may inform the path to rational drug design.

We and others have previously resolved the structures of the SARS-CoV-2 nsp16/nsp10 enzyme complex in the presence of a Cap-0 analogue (^me7^GpppA) and methyl donor SAM (*S*-adenosyl-L-methionine) (Fig. 1a)^8,9^. Here we report two structures of the nsp16/nsp10 heterodimer complex in the presence of a cognate RNA product (Cap-1) that consists of N_1_ and an adjoining N_2_ base (^me7^GpppAmU), and a byproduct of the methylation, SAH (S-adenosyl-L-homocysteine), resolved to 2.3 and 2.5Å, respectively (Fig. 1b-c, S. Table 1). These structures were solved by a molecular replacement method using the previously determined Cap-0 (^me7^GpppA)/SAM structure (PDB ID: 6WKS, hereafter referred to as the “substrate structure”) as a search model^8^. Cap-1 RNA and SAH were unambiguously identified in the difference omit maps (S. Fig. 1 a-c). nsp16 adopts a canonical methyltransferase fold with a central β-sheet flanked by two α-helices on one side and three on the other, similar to the substrate Cap (Cap-0)-bound structure^8^ but with a notable difference as described below.

**Fig. 1.**
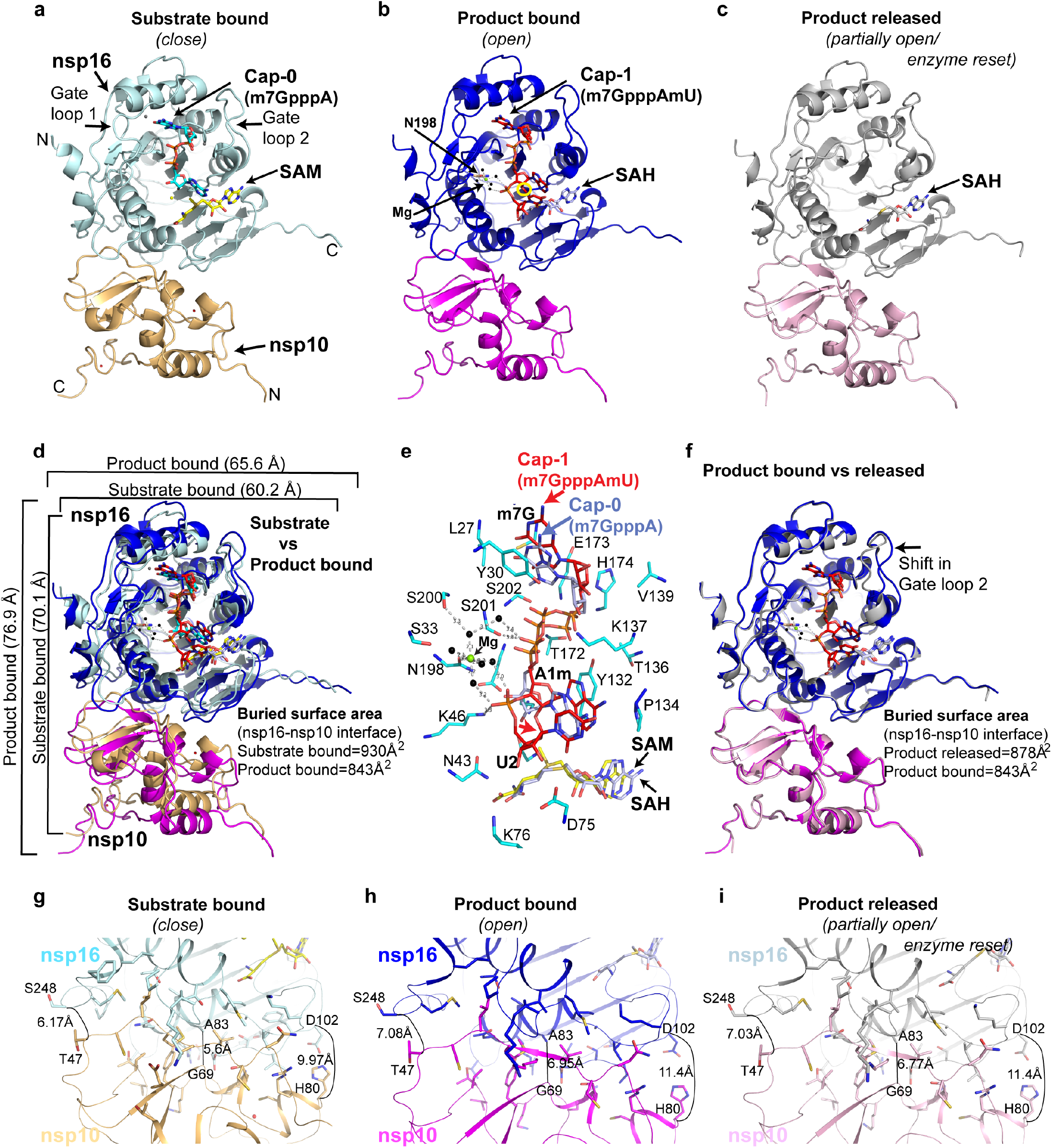
Structures of SARS-CoV-2 nsp16/nsp10 complexes. **a**, The substrate (^me7^GpppA, blue stick) and methyl donor (SAM, yellow stick)-bound nsp16 (cyan)/nsp10 (orange) complex (PDB ID, 6WKS)^8^ represents a closed form. **b**, The product (^me7^GpppAmU, red stick; byproduct SAH [grey stick])-bound nsp16 (blue)/nsp10 (magenta) in an *open* state. A yellow circle shows the methylated ribose (2’-O-me) of N_1_ (A) base. **c**, The SAH (grey) bound nsp16 (grey)/nsp10 (pink) represents a *partially open* or *enzyme reset* state. **d**, Secondary structure-based overlay of nsp16 in substrate-and product-bound states clearly shows the universal expansion of the enzyme upon 2’-O methylation. **e**, A close up view of Cap-binding and catalytic pocket of the product structure shows nsp16 residues (cyan sticks) interacting with Cap-1 (red). A positional change in orientation of the substrate (Cap-0, blue) from the “closed” structure determined previously^8^ is shown. **f**, An overlay of the product (Cap-1) and byproduct (SAH)-bound structures shows change in the orientation of gate loop 2. Reduction in buried surface area between nsp16/nsp10 in *fully* and *partially open* structures (compared to substrate-bound closed state) is shown (**g-i)**.

### nsp16/nsp10 undergoes breathing motion during 2’-O methyl transfer

A structural comparison of nsp16/nsp10 in substrate, product (^m7^GpppAmU and SAH), and byproduct (SAH only) bound structures revealed notable conformational changes in three complexes (Fig. 1). Most strikingly, we observed an overall expansion of the nsp16/nsp10 complex in the product structure as compared to the substrate-bound enzyme (~ 6.8 Å in one dimension and ~ 5.4 Å in the other) although the central β-sheet remains largely unperturbed (Fig. 1d-e, S. Fig. 1i). A positional shift in the Cap-1 analogue also occurs, so that the 2’-*O*-me group on the A1 base now pushes the SAH outward, so that the sulfur atom points away from the 2’-*O*-me moiety, and the carboxy tail of SAH rotates 180° around C_β_ of the SAH (Fig. 1e, S. Fig. 1a-h, j). A comparison of product and byproduct structures reveals no major changes in protein conformations except for an inward shift in gate loop 2 (Fig. 1f). The position of SAH in the product and byproduct structures remains essentially the same, except for the carboxy tail, which rotates back in byproduct structure to assume the original orientation as in the substrate structure (Fig. 1b-d, S. Fig. 1g-h). These changes suggest that the conformation of the SAH-bound enzyme represents a resetting for the next round of catalysis.

The surface area buried (BSA) at the nsp16/nsp10 interface is significantly smaller (843 Å^2^) in the product complex than the substrate structure (930 Å^2^) (Fig. 1d, g). This reduction is less pronounced (BSA=878 Å^2^) in the byproduct structure (Fig. 1f). As a result, the heterodimeric interface in the product structure (and to some extent the byproduct structure) is widened by ~1.5Å at one end, ~ 0.9Å at the other, and ~1.35Å in the center (Fig. 1g-i). Thus, the much-relaxed heterodimeric interface in the product structure appears to be a result of overall widening of the enzyme, which is triggered by a single 2’*-O* methylation event. The enzyme appears to go into a “breathing motion” during catalysis, wherein the substrate, product, and byproduct bound states represent *fully closed, open,* and *partially open (product released/enzyme reset)* states, respectively (Fig. 1). The product RNA Cap, Cap-1 with adjoining uracil as N_2_ base at 3’-end, is accommodated within a deep pocket constituted by one side of the central β-sheet and the two gate loops (gate loop 1 [amino acids 20-40] and gate loop 2 [amino acids 133-143]). The byproduct SAH in both structures binds similarly in a cavity at the C-terminal side of the parallel β-strands except for different orientations of their carboxy tails (S. Fig. 1a-b, g-h).

### Metal dependency for 2’-*O*-methylation by SARS-CoV-2 and its clinical variants

We also observed unambiguous electron density in omit maps at the interface of gate loop 1, a loop between the β8 and β9 strands, and the phosphate moiety of uracil (U_2_), the N_2_ base downstream to the RNA Cap (Fig. 1e and S. Fig. 1a, c). The features of this density suggested a divalent metal ion coordinating with water molecules. Metal ions stabilize nucleic acid substrates in addition to acting as catalytic agents in enzymatic reactions. In other positive single-stranded RNA viruses (e.g., 2’-*O* MTase such as dengue NS5), a magnesium ion stabilizes the RNA cap by coordinating with the inverted triphosphate moiety from the solvent-exposed side of the RNA cap (Fig 2f). A magnesium ion in nsp16 of the previous CoV outbreak strain (PDB ID: 2XYR) binds to a remote site constituted by T58 and S188 located at the opposite face of the catalytic pocket^10^. A direct binding of metals in the substrate/catalytic pocket and their role in 2’-*O* MTase activity of the CoVs nsp16, including SARS-CoV-2, has not been previously suspected.

**Fig. 2.**
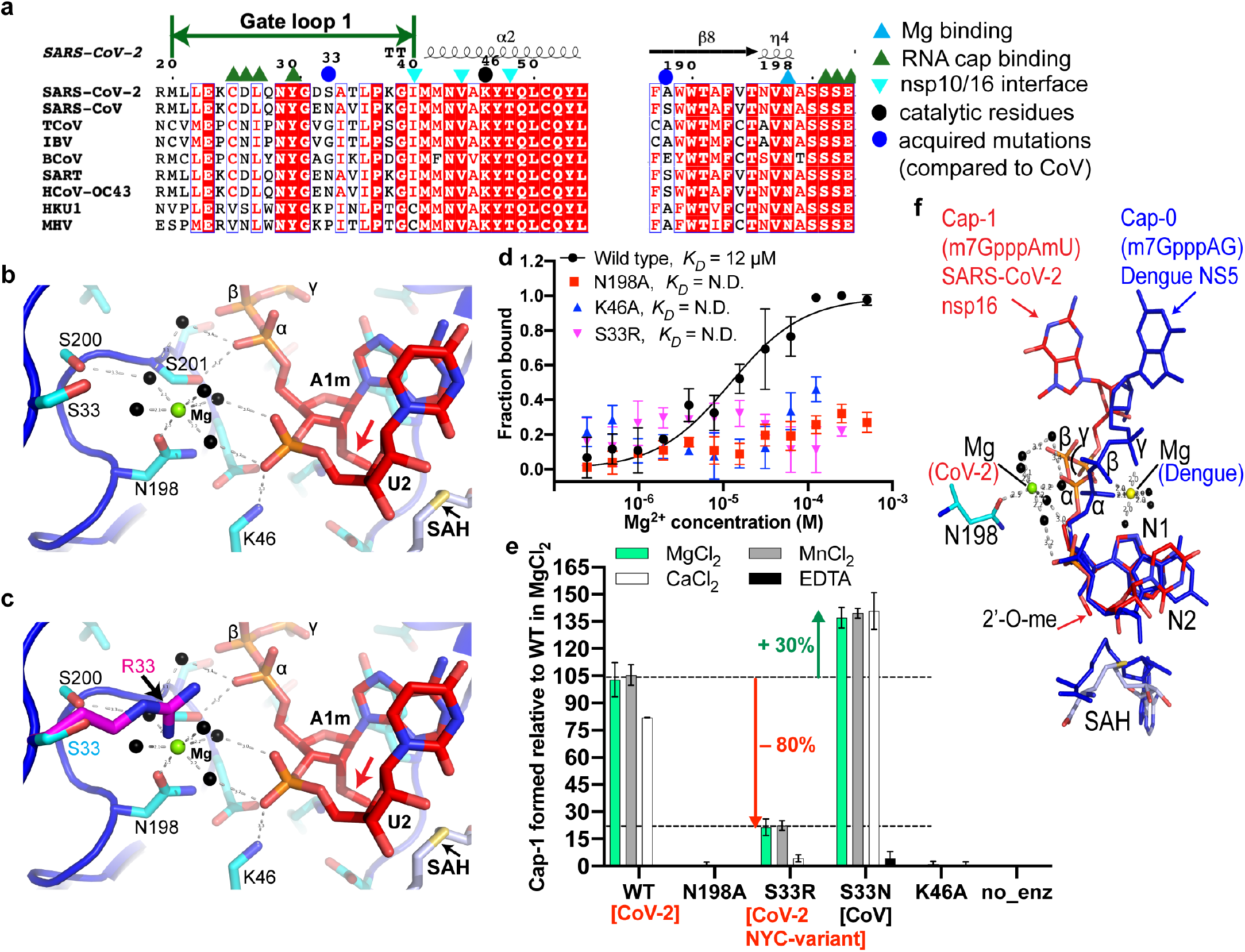
Metal dependency of nsp16/nsp10 and its clinical variant for 2’-*O* methylation., **a**, Alignment of nsp16 from different CoV representing all three sub-classes (α, β, *γ*). Blue sphere denotes S33R, which locates in gate loop 1; S33 is an asparagine (N) in SARS-CoV; S33R is a clinical nsp16 variant of SARS-CoV-2; Cyan triangle, N198 that coordinates Mg^2+^; black sphere, catalytic lysine (K46 of **K**DKE tetrad). **b**, A close-up view of coordination of Mg^2+^ ion (green sphere). Mg^2+^ coordinates with five water molecules (black spheres) at the nucleic acid face and the side chain of N198 on the opposite face. Two of these water molecules hold the phosphoryl oxygens of the U_2_. Red arrow; 2’-*O*-methylated ribosyl of A_1_ base. **c**, The side chain of arginine (magenta, modeled) at the S33 (cyan) position intrudes into the Mg^2+^ pocket and thus, may displace Mg^2+^ or disrupt the Mg^2+^/water network. **d**, Binding isotherms of Mg^2+^ nsp16/nsp10 (WT and mutants) interaction derived from the MST data. (N.D., not determined). **e**, Quantitative measurement of Cap-1 formation by nsp16/nsp10 enzymes experiments (+/- metals, EDTA) as derived from the LC/MS data. Results are averages of three independent experiments (n=3) normalized to a WT dataset consisted of seven datapoints with one standard deviation (s.d.) for each metal or EDTA shown as error bars. Source data are provided as a Source Data File. no_enz, reaction devoid of nsp16/nsp10 enzyme. **f**, An overlay of the N_1_ and N_2_ bases and SAH of SARS-CoV-2 nsp16/nsp10 (red) and dengue NS5 (blue; PDB ID: 5DTO) RNA caps shows entirely different orientations of the terminal base of the cap (^me7^G), two phosphates (β and γ) and Mg^2+^ ions. Mg^2+^ (in dengue, yellow sphere) stabilizes the three phosphates whereas in SARS-CoV-2 (green sphere) it indirectly (water-mediated) stabilizes the phosphate of the N_2_ base on one side and engages N198 of nsp16 on the opposite side. Red arrow; 2’-*O* methyl of ribose of N_1_.

Since our protein purification buffers and crystallization solutions contain magnesium and calcium salts, respectively, we fit these metals into the additional electron density, but could refine the product structure with better statistics when Mg^2+^ was modeled along with its coordinating water molecules (B factor = 52.5 Å^2^ and 73.3 Å^2^ for Mg^2+^ and Ca^2+^, respectively) in this density. As modeled, Mg^2+^ is further stabilized by direct interaction with the side chain of an invariant N198 and assumes a near-ideal octahedral geometry with coordinating water molecules (Fig. 2a-b, S. Fig. 1d-e, j). Consistently, the wild-type (WT), but not the N198A mutant of nsp16/nsp10 binds Mg^2+^ with high affinity (Fig. 2d). A direct interaction of protein to Mg^2+^ and its orientation in the Cap binding pocket is unique to SARS-CoV-2 (Fig. 2c). For example, in dengue NS5, Mg^2+^ is exposed to solvent and cross-links the phosphate groups of the RNA cap without ligating to the protein, whereas it directly binds to nsp16 (through N198) and the phosphate of the U_2_ base (water-mediated) in SARS-CoV-2 (Fig. 2f).

Our previous work revealed the basis of target specificity of nsp16 for adenine nucleotide at the N_1_ position^8^. We and others postulated that K170 (the second lysine of the KD**K**E catalytic tetrad) acts as a general base to facilitate methyl transfer from SAM to 2’-OH of the A1 nucleotide in SARS-CoV-2^8^ and other CoVs^3,10^. The product structure allowed us to examine the roles of K170 and K46 (first lysine of the **K**DKE catalytic tetrad) in detail. The side chains of K170 and K46 form hydrogen bonds with the 2’-*O* of the product (i.e., methylated ribosyl of A1) and phosphoryl oxygen of U_2_ base, respectively (Fig. 2b, S. Fig. 1d-f)). The phosphoryl oxygens of U_2_ also interact with water molecules that coordinate with Mg^2+^. Mutation of K46 and N198 to alanine completely abolished Mg^2+^ binding and catalytic activity of nsp16/nsp10 (Fig. 2d-e). This is consistent with the network of side chains of K46, K170, and N198 that we identify here as important for catalysis. Such an arrangement correctly positions the RNA cap in the catalytic pocket to both ensure efficient 2’-*O* methylation of A1 base and prevent unintended methylation of the adjoining U_2_ base by restricting its movement or misalignment during catalysis of the A1 base. The U_2_ nucleotide is largely exposed to solvent and showed some deviation in the geometry (S. Fig. 1d).

Previously, we mapped the acquired mutations that have been identified in SARS-CoV-2 nsp16 on its structure and postulated their potential roles in RNA binding and/or catalysis^8^. Among these, the S33 residue in gate loop 1 is particularly noteworthy given the high occurrence of the S33R (20755:A>C) mutation in SARS-CoV-2 strain associated in the New York City outbreak^11^, and its mutation to an asparagine (S33N) in a previous CoV outbreak strain (Fig. 2a, S. Fig. 1a). The side chain of S33 resides 6.3Å and 9.6Å away from Mg^2+^ and phosphoryl oxygens of the U_2_ base in the product structure, respectively. The side chain of the arginine in S33R, as we modeled in Fig. 2c, may intrude into this pocket to reduce these distances by ~ 4.18Å, thus disrupting the coordination of Mg^2+^, which would in turn disorient the target base (A1) in the catalytic pocket. A shorter side chain of asparagine would be less intrusive and may provide additional contacts to a divalent metal ion, strengthen RNA binding, and therefore enhance 2’-*O*-methylation.

To further probe these models, we employed LC/MS to measure Cap-1 formation by nsp16/nsp10 WT and variant enzymes, and their dependence on various metal ions. Consistent to magnesium-binding affinity, where mutants N198A, K46A and S33R exhibited negligible Mg^2+^ binding (Fig. 2d), N198 (which directly interacts with magnesium), and K46 (which stabilizes the phosphate of U_2_) when mutated to alanine, completely abolished the enzymatic activity of nsp16/nsp10 (Fig. 2e). Strikingly, the activity of S33R mutant decreased by ~ 80%, further validating our structural interpretation about this clinical variant (Fig. 2c-e). In contrast, the S33N mutation resulted in 30% increased activity, suggesting that the SARS-CoV nsp16, which has N at this position, has superior 2’-*O* methylation capability compared to SARS-CoV-2 nsp16 (Fig. 2e). Moreover, the SARS-CoV-2 nsp16 shows indistinguishable 2’-*O* methyltransferase activity in the presence of magnesium and manganese, but a 20% loss in the presence of calcium. The S33N mutant showed no preference for any of the three divalent ions tested (Mg^2+^, Ca^2+^, Mn^2+^). However, S33R showed comparable activity in the presence of Mg^2+^ and Mn^2+^ (although 80% less than the WT), but residual enzymatic activity in the presence of Ca^2+^ (Fig. 2e).

The unique role of a divalent metal ion in SARS-CoV-2 nsp16 appears to be architectural (Fig. 2f), yet it is essential for accurate and efficient 2’-*O* methylation of the first transcribed base of the SARS-CoV-2 genome. Thus, allowing the virus to evade host innate immune responses. Such reliance and preference for metals also suggests that an imbalance in cellular metal concentrations could differentially alter the RNA capping and thus, host innate immune response to infections by various CoVs. In support of this concept, hypocalcemia is considered as a strong predictor of inhospital COVID-19 deaths^12,13^, and severely ill COVID-19 patients had significantly lower magnesium levels in whole blood^14^. One possibility is that the sub-optimal RNA capping by nsp16/nsp10 variants, together with altered levels of divalent metals, could trigger excessive immune response and cause hypercytokinemia in a subpopulation of COVID-19 positive patients. Future studies should determine the direct correlations between RNA capping, metal levels in the host cellular milieu, and innate immune response.

## METHODS

### Protein expression and purification

The nsp16 (NCBI reference sequence YP_009725311.1) and nsp10 (NCBI reference sequence: YP_0009725306.1) of the seafood market pneumonia SARS-CoV-2 isolate Wuhan-Hu-1 (NC_045512) were cloned into a duet vector, co-expressed in *E. coli,* and purified using the method we previously described ^8^. Briefly, the clarified *E. coli* lysates were loaded on to a metal affinity column and proteins were eluted by running a concentration gradient of imidazole. The N-terminal His6 tag from nsp16 was then proteolytically removed. The tag-free fractions were purified by successive passage through affinity, ion-exchange, and size-exclusion chromatography columns. The purified enzymes were concentrated to 5 mg/mL and used immediately for subsequent biochemical and/or crystallographic studies. We used the same method for all mutant nsp16/nsp10 enzymes reported in this study.

### Crystallization, X-ray diffraction data collection and structure determination

The initial nsp16/nsp10 complex was grown by the sitting drop vapor diffusion in a crystallization solution 10% (v/v) of 2-propanol, 0.1 M MES/NaOH pH 6.0, 0.2 M calcium acetate. After 3-4 rounds of optimization by varying pH, precipitant, and salt concentrations, we grew larger crystals amenable to synchrotron radiation. We soaked these crystals with an RNA analog representative of the Cap-1 structure (^me7^GpppA_(2’-_O_-me)_U). The crystals were cryo-protected by serial soaks in a solution containing the original mother liquor and increasing concentrations (0 to 20% v/v) of ethylene glycol, and then flash-frozen in liquid nitrogen. Crystals of the nsp16/nsp10/Cap-1/SAH and nsp16/nsp10/SAH complexes diffracted X-rays to 2.3 and 2.5 Å resolution with synchrotron radiation, respectively (S. Table 1). Both crystals belong to the space group P3121 with similar unit cell dimensions a=b=184Å, c = 57Å, α = β = 90°, and γ = 120°, and with one nsp16/nsp10 heterodimer per asymmetric unit. All data (measured at wavelength 1.07 Å) were indexed, integrated, and scaled using XDS, aimless, and various ccp4 suite programs integrated into the RAPD pipeline at the NECAT 24ID beamline^15^. The structure was solved by molecular replacement using a Cap-0 ternary complex of nsp16/nsp10/SAM (PDB ID: 6WKS)^8^ structure as a template in Phaser^16^. The resulting maps indicated unambiguous electron densities for RNA Cap-1 and SAH. Ligand topologies and geometrical restraints were generated using PRODRG (http://prodrg1.dyndns.org), GRADE (http://grade.globalphasing.org), and eLBOW (Phenix)^16^ programs. We iteratively rebuilt and refined the model with good stereochemistry using the programs Coot^17^ and Phenix^15^ (Supplementary Table 1). All figures of structural models were generated using Pymol (The PyMOL Molecular Graphics System, Version 2.0 Schrödinger, LLC).

### Determining affinities of WT and mutant nsp16/nsp10 binding to magnesium

We used microscale thermophoresis (MST) to derive equilibrium dissociation constants *(K_D_)* for protein-metal interactions. A detailed protocol has been published^8^. Briefly, 20 μM of each enzyme complex was labeled by incubating with dye solution (60 μM) in the labeling buffer at room temperature for 30 min. The magnesium stock was prepared in the MST reaction buffer (20 mM HEPES pH 7.5, 150 mM NaCl, 0.5% glycerol, and 0.05% Tween 20), and two-fold serial dilutions (from 2 mM stock) were made in 12 steps. The labeled protein (20 nM) was equally mixed into each ligand reaction (ligand concentration ranges 500 nM to 1 mM). The final reaction mixtures were loaded and measured on a Monolith NT.115 instrument (NanoTemper Technologies) at 25 °C. The results shown here are from three independent experiments. Data were fitted by a singlesite binding model in GraphPad Prism (GraphPad Software, San Diego, CA).

### Enzyme activity assay

We used a LC/MS-based method (reported earlier)^8^ for quantitative measurement of Cap-1 formation by nsp16/nsp10 enzymes in the presence of various metals. Briefly, 0.1 μM of enzymes were allowed to react with 1 μM ^me7^GpppA-capped 25 nt RNA in a buffer (50 mM Tris-HCl, pH 8.0, 5 mM KCl, 1 mM DTT, 0.2 mM SAM) supplemented with 1 mM MgCl_2_ or MnCl_2_ or CaCl_2_ or 5 mM EDTA. The reactions were incubated at 37°C for 30 min, stopped by heating at 75°C for 5 min in the presence of 5 mM EDTA, and were subjected to LC/MS intact mass analysis. Nucleic acids in the samples were separated using a Thermo DNAPac™ RP Column on a Vanquish Horizon UHPLC System, followed by mass determination using a Thermo Q-Exactive Plus mass spectrometer. The raw data were deconvoluted using Promass HR (Novatia, LLC). The deconvoluted mass peak ratios between reactants and the expected products was used to estimate the percentage of 2’-*O* methylation. Results shown in Fig. 2e are average of three independent experiments (n=3) normalized to a WT dataset consisted of seven data points. Source data are provided as a Source Data File. The RNA substrate used in this assay is ^me7^GpppAUAGAACUUCGUCGAGUACGCUCAA-[6-FAM].

## DATA AVAILABILITY

The information about coding sequences of nsp16 (NCBI reference sequence YP_009725311.1) and nsp10 (NCBI reference sequence: YP_0009725306.1) of the seafood market pneumonia SARS-CoV-2 isolate Wuhan-Hu-1 (NC_045512) used in this study is available at NCBI (https://www.ncbi.nlm.nih.gov/nuccore/NC_045512). Files for atomic coordinates and structure factors were deposited in the Protein Data Bank under accession codes 7LW3 (product bound) and 7LW4 (SAH bound). Correspondence and requests for material should be addressed to Y.K.G.

## Acknowledgments

This work was partially supported by funding from the San Antonio Partnership for Precision Therapeutics (SA-PPT), pilot awards from the Institute for Integration of Medicine and Science (IIMS) and UT Health San Antonio (UTHSA), and the Max and Minnie Tomerlin Voelcker Foundation to Y.K.G. T.V. is supported by a research training award (RP170345) from the Cancer Prevention Research Institute of Texas (CPRIT). Y.K.G. is also supported by a high impact/high risk award from the CPRIT, and a Rising STARs award from the UT System. We are grateful to beamline scientists at NECAT-24ID sector, APS, Chicago for provision of synchrotron beamtime and data collection. This work is based on research conducted at the Northeastern Collaborative Access Team beamlines (NIH P30GM124165, and DOE DE-AC02-06CH11357 grants). We also thank the X-ray Core of UTHSA.

## Author contributions

Y.K.G. conceived, designed, and supervised the overall study, and performed crystallographic studies; T.V., A.M., S.A. purified proteins, T.V. performed crystallization, S.Q. and T.V. performed biochemical assays. S.-H.C. performed the LC/MS-based assays with assistance from N.D. L.M.-S. provided reagents. Y.K.G. wrote the manuscript, and all authors have read and approved this version.

## Materials and correspondence

Files for atomic coordinates and structure factors were deposited in the Protein Data Bank under accession codes 7LW3 (product bound) and 7LW4 (SAH bound). Correspondence and requests for material should be addressed to Y.K.G.

## SUPPLEMENTARY INFORMATION

**S. Fig. 1.**
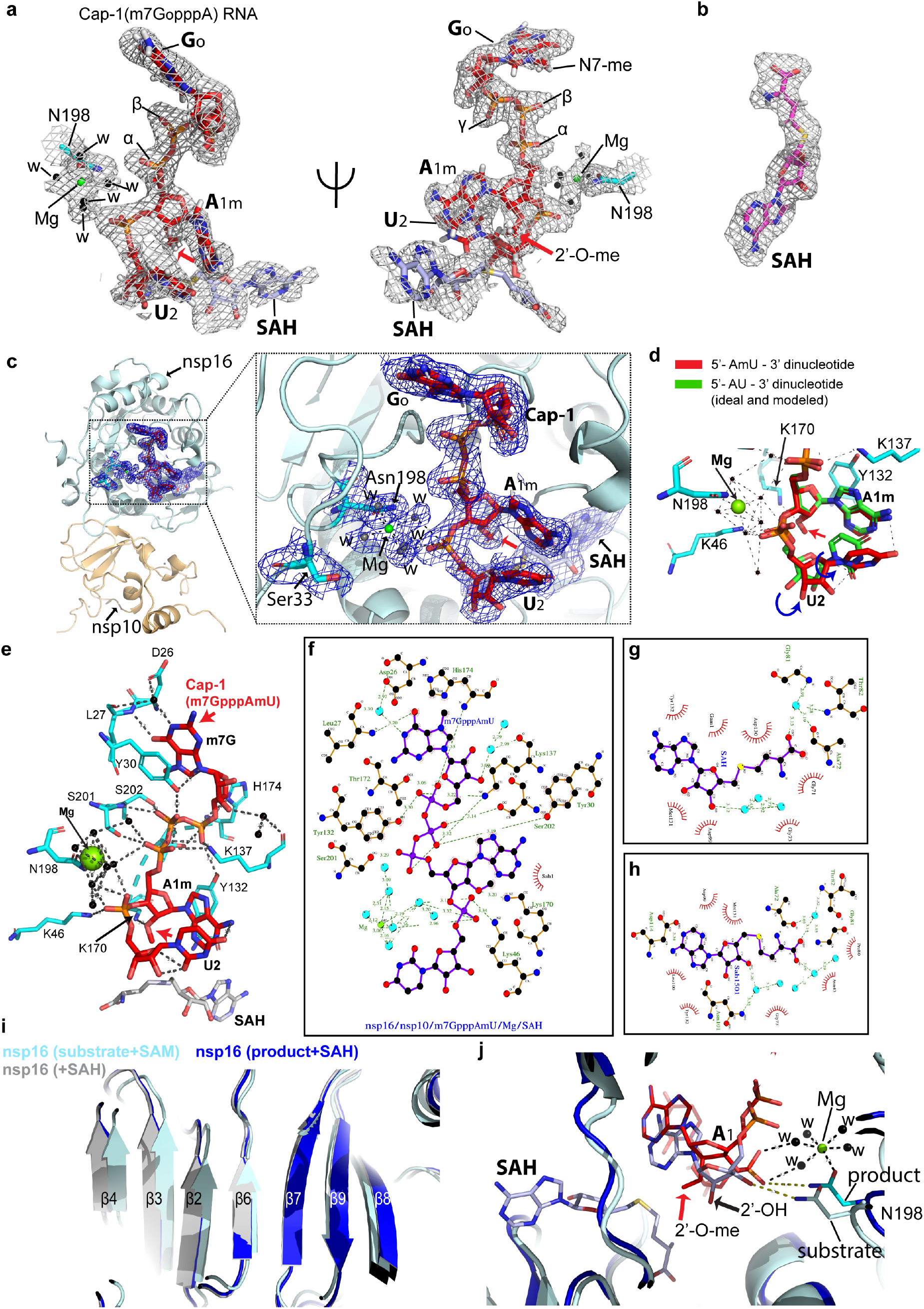
**a**, A close-up view of the product (^m7^GpppAmU, red stick), SAH (grey stick), magnesium (green sphere), water (W, black spheres), and N198 binding. The Fo-Fc electron density omit map for these ligands contoured at 2.7σ is shown as a grey mesh. To minimize the possibility of bias introduced by ligands, all ligands were excluded from refinement and phase calculations. Positions of the methyl group attached to the acceptor moiety (2’-O in ribose of the target nucleotide A1m) is depicted by red arrow. **b**, The Fo-Fc electron density omit map for SAH (red stick) in nsp16/nsp10/SAH structure contoured at 2.7σ is shown as a grey mesh. **c**, Final structure of nsp16 (cyan)/nsp10(orange)/^me7^GopppAmU (red)/SAH (grey)/ Mg^2+^ (green)/Mg^2+^-coordinating waters showing 2Fo-Fc map (blue mesh) contoured at 1.0σ. **d**, An overlay of an A in an ideal AU (green) dinucleotide over Am base in the product structure shows deviated geometry (outward motion, blue arrows) of the U_2_ base, suggestive of a state preceding the product’s release. **e**, nsp16 residues (cyan) that directly interact with Cap-1 RNA (red) and Mg^2+^ (green sphere) and water (black sphere). **f**, Protein-ligand interaction network of nsp16/nsp10/Cap-1 (^m7^GpppAmU) and SAH in product (**g**), and SAH-bound (**h**) nsp16/nsp10 structures. The green dashed lines represent hydrogen bonding, and the cyan spheres represent water molecules. These figures were generated using the LigPlot+ program^1^. **i**, A secondary structure-based superposition of nsp16 in all three structures show good overlay of the central β-sheet in all three structures. The regions flanking the central core in nsp16 and the entire nsp10 universally expands (relative to substrate/SAM-bound form, cyan) to assume a more relaxed (or fully open state) in the product-bound form (blue). The structures of product plus SAH and only SAH-bound enzymes do not deviate except in the gate loop region. **j**, An overlay of the substrate (light cyan) and product (blue) structure is shown with their respective Caps (Cap1U as red stick in product, grey stick; Cap-0 in substrate structures). In the absence of Mg^2+^ in the substrate structure, the side chain of N198 coordinates with 3’-OH of the N_1_ base whereas in the product structure it coordinates with the phosphoryl oxygen of the N_2_ base and Mg^2+^ ion.

**Supplementary Table 1.**
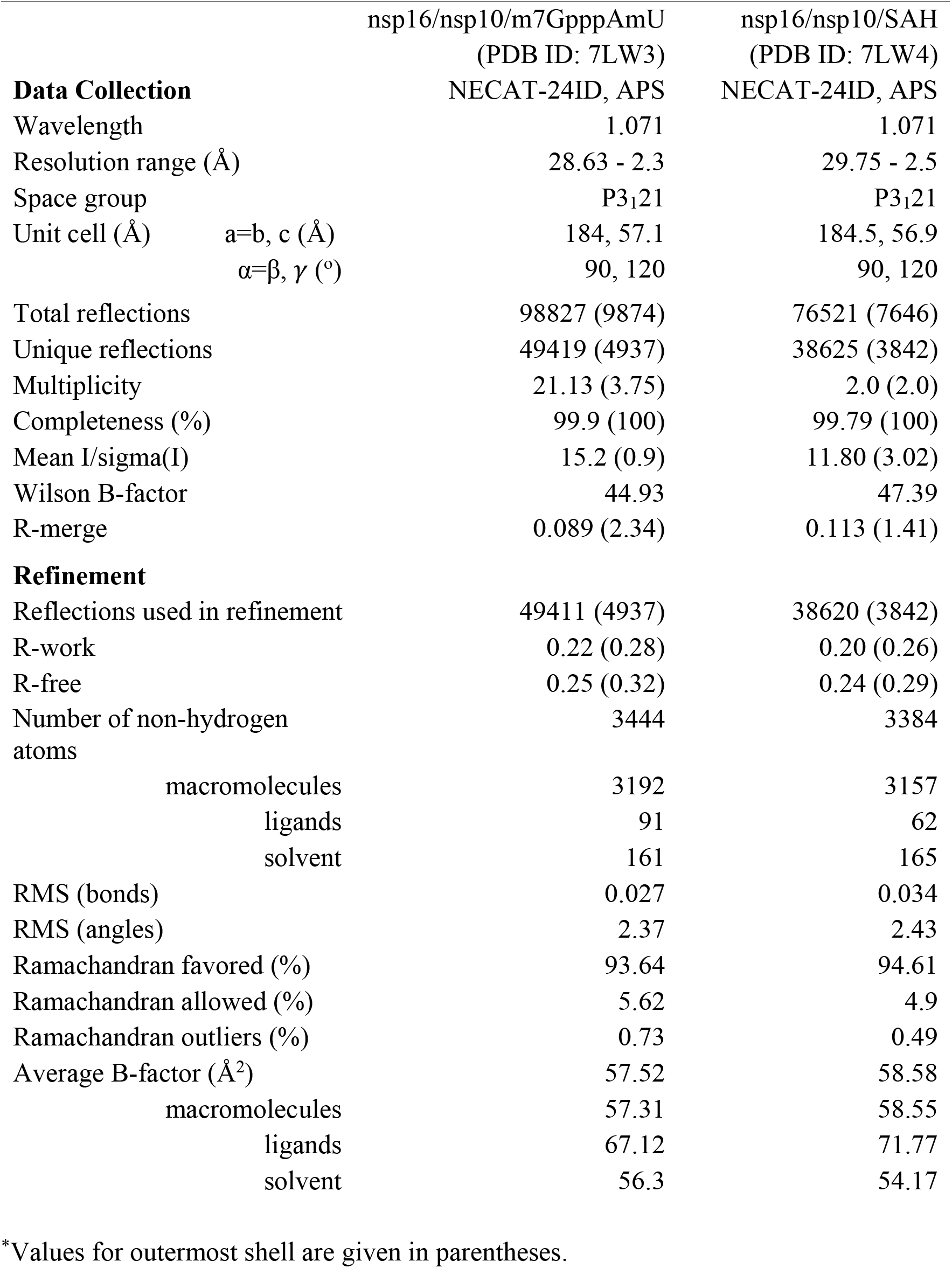
Data collection and refinement statistics (molecular replacement)

